# Sialoglycoconjugate Profiling of Human Choroid, Retinal Pigment Epithelium, and Macular Degeneration Related Lesions

**DOI:** 10.1101/2025.04.23.650306

**Authors:** Emma M. Navratil, Piper A. Wenzel, Miles J. Flamme-Wiese, Jack E. B. Miller, Luke A. Wiley, Edwin M. Stone, Budd A. Tucker, Robert F. Mullins

## Abstract

Age-related macular degeneration is a leading cause of central vision loss in the elderly. Early hallmarks of the disease include basal laminar deposit and choriocapillaris degeneration. The location and composition of sialoglycoconjugates in healthy and diseased choroid and disease-related lesions have not been thoroughly examined. This study utilized lectins to examine sialoglycoconjugates in human tissue, specifically *Sambucus nigra*/Elderberry Bark Lectin (EBL) and *Maackia amurensis* lectin II (MAL-II), to examine α-2,6 and α-2,3 sialic acids, respectively. EBL and MAL-II both label the choroid and basal laminar deposit, with slightly different patterns. Whereas MAL-II predominantly labels the choriocapillaris endothelium, EBL also labels Bruch’s membrane and extracellular domains surrounding the vasculature (intercapillary pillars). EBL labeling overlaps with the distribution of complement factor H to a greater extent than MAL-II. After treatment with neuraminidase to remove terminal sialic acids, a battery of lectins was applied to sections of choroids. Lectins that recognize β-galactose, N-acetyllactosamine, galactose (β-1,3) N-acetylgalactosamine, and α - or β-N-acetylgalactosamine showed increased reactivity, including increased labeling of glycans in basal laminar deposits. This study provides insight into the location and partial identities of sialoglycoconjugates in the human choroid, with possible implications for the pathogenesis of macular degeneration.

## INTRODUCTION

Age-related macular degeneration (AMD) is a leading cause of central vision loss in older adults. In adults over 40, early AMD has been reported to have a prevalence of 11.64% and late AMD has a prevalence of 1.49% (1). Early in the development of AMD, the choriocapillaris (the microvasculature that supplies the photoreceptor cells) begins to degenerate below the retinal pigment epithelium (RPE), decreasing choroidal blood flow and nutrient availability to the RPE and overlying retina. Pathologic deposits called drusen, basal laminar deposits, and basal linear deposits appear between Bruch’s membrane and the RPE. The RPE becomes dysfunctional and can also degenerate.

Late-stage AMD exists in two manifestations: neovascular (wet) and geographic atrophic (dry), which can exist independently or in tandem. In choroidal neovascular AMD, vessels from the choroid breach Bruch’s membrane and grow into the sub-RPE and/or subretinal space, disrupting the ionic composition necessary for vision. In some cases, pathologic vessels arise from the retinal circulation. In geographic atrophy, the choroid, RPE, and photoreceptors die, leaving patches of degeneration in the macula.

Risk factors for AMD include age, smoking, and several genetic loci, several of which are in complement regulatory genes such as Y402H in *CFH* (rs1061170) (2–4). Homozygosity for the Y402H risk allele of *CFH* is associated with increased complement activation and deposition of the membrane attack complex (MAC) of complement in the choriocapillaris (5). Prevention and treatment options for AMD are limited. A special formulation of vitamins (AREDS2) has shown efficacy in slowing the progression of intermediate AMD to late AMD (6). Lifestyle changes such as adopting a Mediterranean diet (7–10), exercise (11, 12), and ceasing smoking (13–15) can help decrease the risk of AMD development or progression. Neovascularization in late AMD is treated with intravitreal injections of anti-VEGF (vascular endothelial growth factor) drugs. More recently, the complement inhibitors pegcetacoplan and avacincaptad pegol have received FDA approval for treatment of geographic atrophy, though these drugs increase the occurrence of neovascularization (16, 17). Overall, there is significant progress to make in terms of treating and preventing this disease. A better understanding of the complement system and its interactions with ocular cells is warranted.

Complement inhibitors such as factor H (FH) bind cells and the extracellular matrix in part through protein-carbohydrate interactions. Cells are coated with a layer of glycoconjugates which are often capped with sialic acids: nine-carbon monosaccharides derived from neuraminic acid that have a keto acid functional group (18). More than 50 sialic acid species exist, with N-acetylneuraminic acid (Neu5Ac) and N-glycolylneuraminic acid (Neu5Gc) being the most common sialic acids found in most mammals. In humans, Neu5Ac is the most common and Neu5Gc is absent due to a mutation in the gene encoding CMP-NeuAc hydroxylase that converts Neu5Ac to Neu5Gc in other species (19, 20). Sialic acids are typically found at the terminals of glycan chains. The process of sialylation occurs via α-2,3 and α-2,6 linkages of sialic acid moieties to galactose or N-acetylgalactosamine on O-linked and N-linked glycan chains. Sialic acids can also exist as polysialic acid chains through α −2,8 and α −2,9 linkages. A variety of sialyltransferases exist with different donor/acceptor specificities and binding conformations they can produce (21).

In addition to its interactions with the complement system, the glycan coating of cells informs the way cells interact with other cells and the extracellular matrix. Sialic acids are anti-adhesive due to the negative charge they confer to the cell surface. Vascular endothelial cells and erythrocytes both have a high concentration of sialylated glycans on the cell surface, which causes repulsion between the cells and allows erythrocytes to circulate unimpeded by attachment to the vessel walls or other blood cells (22, 23). Immune cells commonly possess sialic acid receptors including sialic acid-binding immunoglobulin-like lectins (SIGLECs) and selectins, which contribute to modulation of immune signaling and endocytosis (24).

Despite these known essential functions of glycoconjugates and sialic acids, investigation of glycoconjugates in relation to aging and AMD is limited. Soft and hard drusen contain glycoconjugates that are bound by the lectinsConA, LCA, LFA, RCA-120, and WGA; this indicates the presence of mannose, glucosamine, galactose, and sialic acid (25). Drusen possess cores that bind to PNA after neuraminidase treatment and do not contain lipids, suggesting that the drusen cores contain glycoproteins with O-linked carbohydrate chains (26). Lectin labeling of choroidal neovascular membranes (CNVMs) showed different labeling intensities between normal and pathological vessels. Specifically, SBA and sWGA (which bind α −/β-linked GalNAc and GlcNAc respectively) label endothelial cells of CNVMs (27). Many proteins that have specific roles in AMD pathogenesis are glycoproteins, including vascular endothelial growth factor (VEGF), the sialomucin CD34 (which shows age-related decline in the choriocapillaris (28)), intercellular adhesion molecule 1 (ICAM1), complement factor H (FH), factor H-like protein 1 (FHL1), and the complement factor H related proteins 1-5 (CFHR1-5). Kliffen and colleagues investigated glycosaminoglycans in maculae with and without AMD, finding chondroitin 4-sulfate and heparan sulfate in basal laminar deposits and hyaluronic acid in nodular drusen, confluent drusen, and intercapillary pillars. Maculae with AMD also had higher amounts of total glycosaminoglycans than control maculae (29). AMD related pathologic basal laminar deposits contain ɑ-D-GalNAc, which is not present in other macular structures (30). Recently, Swan et. al. performed elegant biochemical studies characterizing the sialome of the retina in relation to FH binding (31).

Collectively, this evidence suggests that glycoconjugates and specifically sialic acids could play a role in AMD pathogenesis. In this study we used lectins, antibodies, and microscopy to examine sialic acids and their underlying carbohydrate moieties in healthy RPE and choroid, as well as in eyes with basal laminar deposit and choroidal neovascular membranes.

## METHODS

### Tissue Acquisition and Preparation

Human eyes (n=10) were collected by the Iowa Lions Eye Bank with full consent from the next of kin for use in research and in compliance with the Declaration of Helsinki. An 8-12mm punch centered on the macula was collected from each eye and fixed in 4% PFA, cryopreserved in sucrose solution, and embedded in optimal cutting compound (Tissue-Tek O.C.T., Sakura Finetek #4583, Torrance, CA). 7 µm thick cryosections were obtained for epifluorescence analysis. 12-16 µm thick sections were obtained for confocal microscopy analysis.

### Tissue Treatments

For experiments characterizing the chemical identities of sialic acid-containing glycoconjugates, sections were treated with 2:1 chloroform:methanol by placing the slides at an angle in a glass dish and repeatedly pipetting the solution over the section (approximately 6 times). Extraction of lipids with chloroform and methanol is derived from the Folch Method (32) and adjusted for use on tissue sections as opposed to tissue homogenate (33). Lipid removal by this treatment was confirmed with 1% w/v Sudan Black B (Thermo Fisher Scientific #AC419830100, Waltham, MA) in 70% ethanol. Sections were incubated for 5 minutes with Sudan Black B solution, followed by destaining with 70% ethanol. Sections were compared to non-extracted controls on adjacent sections.

To identify the penultimate carbohydrate epitopes in the aging eye masked by sialic acids, sections were incubated with α −2-3,6,8 neuraminidase in 1x GlycoBuffer or 1x GlycoBuffer only (NEB #P0720L, Ipswich, MA) for 16 hours at 37C. Enzyme treated sections received 200 units of neuraminidase each. Lectin histochemistry was conducted immediately following enzyme incubation. Neuraminidase treated sections were compared to buffer only controls on adjacent sections.

### Lectin Histochemistry

All lectin histochemistry steps were conducted in PBS with 1mM MgCl2 and CaCl2 divalent cations. Sections were blocked with 0.1% w/v BSA (RPI #A30075, Mt. Prospect, IL) for 15 minutes. Sections were then incubated with the biotinylated lectins at the dilutions listed in Table 1 for 30 minutes, followed by 3 5-minute washes. Following washes, sections were incubated for 30 minutes in 25 µg/mL avidin-Texas red (Vector Laboratories #A-2006-5, Newark, CA) and 0.2 µg/mL DAPI (Thermo Fisher Scientific #D21490, Waltham, MA). Another set of 3 5-minute washes was conducted, and slides were then cover slipped in Aqua-Mount (Epredia #13800, Kalamazoo, MI). Lectins, sources, and working concentrations are listed in Table 1.

**Table 1.** Lectin information. All lectins were obtained from Vector Labs, Newark, CA.

| Abbreviation | Source | Binding Moiety | Working Concentration | Catalog # |
| --- | --- | --- | --- | --- |
| AAL | *Aleuria aurantia* | α-Fucose | 40 μg/mL | B-1395 |
| ABL | *Agaricus bisporus* | GlcNAc | 40 μg/mL | B-1425 |
| ACL | *Amaranthus caudatus* | Gal-β-3GalNAc | 40 μg/mL | B-1255 |
| BanLec | *Musa paradisiaca* | Mannose | 40 μg/mL | B-1415 |
| BPL | *Bauhinia purpurea* | Terminal β-galactose | 40 μg/mL | B-1285 |
| ConA | *Canavalia ensiformis* | Mannose | 100 μg/mL | B-1005 |
| DBA | *Dolichos biflorus* | Forssman antigen | 100 μg/mL | B-1035 |
| DSL | *Datura stramonium* | Poly-LacNAc | 40 µg/mL | B-1185 |
| EBL (SNA) | *Sambucus nigra* | α-2,6 sialic acid | 40 µg/mL | B-1305 |
| ECL | *Erythrina cristagalli* | Terminal type 2<br>LacNAc | 12.5 µg/mL | B-1145 |
| EEL | *Euonymus*<br>*europaeus* | B & H blood groups,<br>galactosyl (α-1,3)<br>galactose | 40 µg/mL | B-1335 |
| GNL | *Galanthus nivalis* | Terminal mannose α1-3, α1-6 | 40 µg/mL | B-1245 |
| GSL-I | *Griffonia*<br>*simplicifolia* | Terminal α-galactose | 10 µg/mL | B-1205 |
| GSL-II | *Griffonia*<br>*simplicifolia* | Terminal GlcNAcβ | 40 µg/mL | B-1215 |
| Jacalin | *Artocarpus*<br>*integrifolia* | O-linked<br>oligosaccharides,<br>prefers galactosyl (β-1,3) GalNAc | 100 µg/mL | B-1155 |
| LCA | *Lens culinaris* | α-mannose | 100 µg/mL | B-1045 |
| LEL | *Lycopersicon*<br>*esculentum* | Type 2 poly-LacNAc | 40 µg/mL | B-1175 |
| LTL | *Lotus*<br>*tetragonolobus,* | Fucose α-1,3 | 40 µg/mL | B-1325 |
|  | *Tetragonolobus purpurea* |  |  |  |
| MAL-I | *Maackia amurensis* | Gal (β-1,4) GlcNAc | 40 μg/mL | B-1315 |
| MAL-II | *Maackia amurensis* | α-2,3 sialic acid | 10 μg/mL | B-1265 |
| MPL | *Maclura Pomifera* | Core 1 & 3 O-glycans | 40 μg/mL | B-1345 |
| PHA-E | *Phaseolus vulgaris* | Biantennary<br>galactosylated<br>GlcNAc | 40 μg/mL | B-1125 |
| PHA-L | *Phaseolus vulgaris* | β1-6 branched N-glycans | 40 μg/mL | B-1115 |
| PNA | *Arachis hypogaea* | Gal β-1,3 GalNAc | 100 μg/mL | B-1075 |
| PSA | *Pisum sativum* | Core Fucose | 100 μg/mL | B-1055 |
| PTL-I | *Psophocarpus tetragonolobus* | Blood groups A & B | 40 μg/mL | B-1365 |
| PTL-II | *Psophocarpus tetragonolobus* | Type 2 blood group H | 40 μg/mL | B-1405 |
| RCA120 | *Ricinus communis* | Terminal Type 2<br>LacNAc | 100 μg/mL | B-1085 |
| SBA | *Glycine max* | Terminal $\alpha$ -GalNAc<br>and β-GalNAc,<br>Forssman antigen | 12.5 μg/mL | B-1015 |
| sConA | *Canavalia ensiformis* | α-mannose, α-glucose | 100 μg/mL | B-1005S |
| SJA | *Sophora japonica* | Terminal poly-type 2<br>LacNAc | 40 μg/mL | B-1135 |
| STL | *Solanum tuberosum* | Internal Type 2<br>LacNAc | 40 μg/mL | B-1165 |
| sWGA | *Triticum vulgaris* | GlcNAc | 12.5 μg/mL | B-1025S |
| UEA-I | *Ulex europaeus* | α-linked fucose | 40 μg/mL | B-1065 |
| VVA | *Vicia villosa* | Terminal LacdiNAc,<br>terminal β-GalNAc | 5 μg/mL | B-1235 |
| WFA | *Wisteria floribunda* | Terminal α-GalNAc<br>and β-GalNAc | 40 μg/mL | B-1355 |
| WGA | *Triticum vulgaris* | Terminal β-GlcNAc | 100 μg/mL | B-1025 |

### Immunofluorescence

For some experiments, lectin labeling was colocalized with antibodies directed against cellular or extracellular epitopes. Two sets of colabeling were performed, with blocking, washes, and coverslipping conducted as described above. In the first set, sections were incubated with 4 µg/mL anti-CD34 (Thermo Fisher Scientific #PIMA110202, Waltham, MA), 2.5 µg/mL anti-Collagen IV (Millipore Sigma #AB748, Burlington, MA), and either *Sambucus nigra*/Elderberry bark Lectin (EBL) or *Maackia amurensis* lectin II (MAL-II) for 90 minutes. After washes, sections were incubated with 25 µg/mL avidin-Texas red, 2.5 µg/mL donkey anti-rabbit IgG Alexa Fluor 647 (Invitrogen #A-31573, Waltham, MA), 2.5 µg/mL donkey anti-mouse IgG Alexa Fluor 488 (Invitrogen #A-21202, Waltham, MA), and 0.2 µg/mL DAPI for 45 minutes. For the second set, sections were first incubated with 25 µg/mL anti-FH (R&D Systems #MAB4779, Minneapolis, MN) for 90 minutes, followed by a donkey anti-mouse IgG Alexa Fluor 488 fab fragment antibody (Jackson ImmunoResearch #715-547-003, West Grove, PA) for 45 minutes. Sections were next incubated with 0.15 µg/mL anti-C5b-9 (Agilent #M077701-5, Santa Clara, CA) and EBL or MAL-II for 90 minutes, followed by 10 µg/mL streptavidin DyLight 549 (Vector Laboratories #SA-5549-1, Newark, CA), donkey anti-mouse IgG Alexa Fluor 647 (Invitrogen #A-31571, Waltham, MA), and 0.2 µg/mL DAPI for 45 minutes. Sections were washed between each of the four antibody incubations.

### Epifluorescence Imaging

Epifluorescent images were captured using a Spot RT3 Color Slider (Spot Imaging Solutions RT2540, Sterling Heights, Michigan) attached to an Olympus BX41TF upright fluorescence microscope (Olympus, Tokyo, Japan). Images were scored for the following anatomical components for each lectin and each donor, with and without neuraminidase: apical RPE, basolateral RPE, Bruch’s membrane, choriocapillaris, larger choroidal blood vessels, stromal cells, stromal matrix, BLamD, and choroidal neovascular membranes. Scores were defined as follows: no labeling (0), weak labeling (1), medium labeling (2), and strong/maximum labeling (3). To determine the changes after neuraminidase treatment displayed in Table 1, the score with neuraminidase was subtracted from the corresponding score without neuraminidase, for each donor/lectin/component combination. The score change for each lectin/component combination was averaged for all donors. The changes are displayed in Table 2.

**Table 2.** Lectin histochemistry of the RPE/choroid and AMD lesions with neuraminidase treatment. The change in labeling intensity from control to neuraminidase treatment in different RPE/choroid compartments, basal laminar deposits, and CNV is shown. Changes were categorized as strong increase (+++), moderate increase (++), weak increase (+), no change (0), weak decrease (-), moderate decrease (--), or strong decrease (---). RPE, retinal pigment epithelium; BM, Bruch’s membrane; CC, choriocapillaris; large blood vessels, LBV; stromal matrix, SM; stromal cells, SC; basal laminar deposit, BLamD, choroidal neovascular membranes, CNVMs.

| **Lectin** | **Basal RPE** | **Apical RPE** | **BM** | **CC** | **LBV** | **SM** | **SC** | **BLamD** | **CNVMs** |
| --- | --- | --- | --- | --- | --- | --- | --- | --- | --- |
| AAL | 0 | 0 | + | + | + | + | + | + |  |
| ABL | 0 | 0 | 0 | + | 0 | + | + | 0 |  |
| ACL | 0 | + | 0 | ++ | 0 | + | + | + |  |
| BanLec | 0 | + | 0 | 0 | 0 | 0 | 0 | 0 |  |
| BPL | + | ++ | + | ++ | ++ | + | + | +++ | + |
| ConA | 0 | - | 0 | 0 | 0 | - | - | + |  |
| DBA | 0 | 0 | 0 | 0 | 0 | 0 | 0 | 0 |  |
| DSL | 0 | 0 | + | + | + | + | + | + |  |
| EBL | - | -- | - | -- | -- | -- | -- | --- | -- |
| ECL | ++ | ++ | + | ++ | + | + | + | +++ | - |
| EEL | 0 | + | + | ++ | + | + | + | + |  |
| GNL | 0 | + | + | 0 | + | - | 0 | ++ |  |
| GSL-I | 0 | 0 | 0 | 0 | 0 | 0 | 0 | 0 |  |
| GSL-II | 0 | 0 | 0 | 0 | 0 | 0 | 0 | 0 |  |
| Jacalin | 0 | 0 | + | - | - | + | + | + |  |
| LCA | 0 | 0 | 0 | 0 | + | + | + | 0 |  |
| LEL | - | - | 0 | 0 | - | - | - | 0 |  |
| LTL |  | 0 | 0 | 0 | 0 | 0 | 0 | 0 |  |
| MAL-I | -- | -- | - | -- | -- | -- | -- | - |  |
| MAL-II | - | -- | 0 | -- | -- | -- | -- | - |  |
| MPL | 0 | 0 | 0 | 0 | 0 | 0 | 0 | + |  |
| PHA-E | - | - | 0 | + | 0 | 0 | 0 | 0 |  |
| PHA-L | + | + | 0 | + | + | + | + | + | 0 |
| PNA | ++ | ++ | + | +++ | +++ | ++ | ++ | ++ |  |
| PSA | 0 | + | 0 | + | + | + | + | + |  |
| PTL-I | 0 | 0 | 0 | 0 | 0 | 0 | 0 | 0 |  |
| PTL-II | 0 | 0 | 0 | 0 | + | 0 | 0 | + |  |
| RCA120 | + | + | + | + | 0 | 0 | - | + | + |
| SBA | ++ | + | + | ++ | ++ | + | ++ | +++ | ++ |
| sConA | + | + | 0 | + | + | + | + | + |  |
| SJA | + | ++ | + | ++ | ++ | ++ | + | + | ++ |
| STL | 0 | 0 | + | 0 | - | - | 0 | 0 |  |
| sWGA | 0 | - | 0 | 0 | - | 0 | - | + |  |
| UEA-I | 0 | 0 | 0 | 0 | 0 | 0 | 0 | 0 | 0 |
| VVA | 0 | 0 | 0 | ++ | + | + | + | ++ |  |
| WFA | ++ | ++ | ++ | ++ | ++ | ++ | ++ | +++ | ++ |
| WGA | - | - | - | - | - | - | - | - |  |

### Confocal Imaging

For colocalization of different sialic acid moieties at higher resolution, sections were imaged using a Leica TCS SPE upright confocal microscope system (Leica Microsystems, Wetzlar, Germany). The step size was set to 0.5 µm. Images were pseudo colored using Leica LAS X software, with far-red channel images shown in blue and UV channel images shown in gray. To generate figures, ImageJ was used to split single color images into RGB color channels via Image > Color > Split Channels. This generated grayscale images, and the image corresponding to the original color was used as depicted in each figure.

## RESULTS

Lectin histochemistry was performed using two lectins, EBL and MAL-II, that recognize sialic acids. EBL has a preference for α-2,6 sialic acids, while MAL-II preferentially binds α-2,3 sialic acids. Aged control and early AMD maculae were evaluated for EBL and MAL-II labeling patterns. In the control choroid, EBL labels the apical and basolateral surfaces of the RPE, as well as Bruch’s membrane (Fig. 1). There is also strong labeling of vessel walls, including capillaries and larger vessels. EBL further recognizes ligands in the intercapillary pillars, which are dense extracellular matrix domains between individual capillaries (Fig. 1). Weaker, diffuse, mottled labeling is also present in the choroidal stroma and sclera. MAL-II labels the apical RPE and photoreceptor outer segments, but not basolateral RPE or Bruch’s membrane. It also labels blood vessel walls, and, unlike EBL, does not consistently react with intercapillary pillars (Fig. 1). Modest stromal labeling appears weaker than EBL, though similar in pattern.

**Figure 1.**
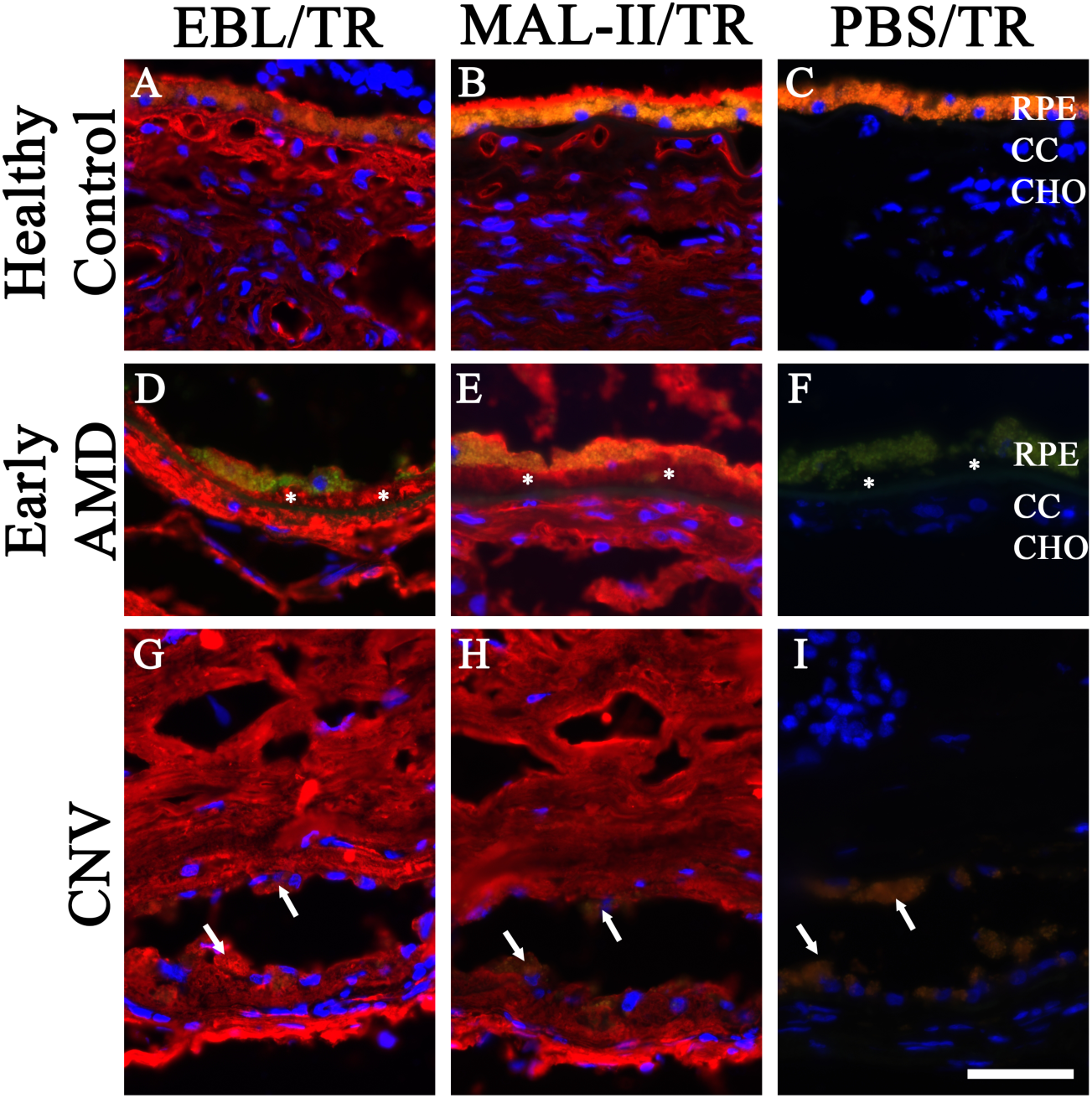
Labeling patterns of sialic acid binding lectins in human choroid. Fluorescence micrographs of human donor choroids including a healthy control (A-C), early AMD (* = BLamD) (D-F), and choroidal neovascular membrane (CNV) (G-I). Choroids were labeled with DAPI and EBL (A,D,G), MAL-II (B,E,H), or DAPI and Avidin Texas Red (TR) only (C,F,I). EBL and MAL-II are shown in red, DAPI is in blue. RPE, retinal pigment epithelium; CC, choriocapillaris; CHO, outer choroid. Yellow-orange autofluorescence in control panels is due to RPE lipofuscin. Arrows indicate RPE cells in the CNV panels. Scale bar is 50 µm.

Lectin labeling was also observed in eyes with basal laminar deposit (BlamD). EBL displays intense, patchy labeling in the deposit itself, while MAL-II shows modest diffuse labeling (Fig. 1, asterisks). In eyes with choroidal neovascular membranes and layers of subretinal fibrosis, the EBL and MAL-II labeling appears similar to labeling of the choroidal stroma in the control donor, with MAL-II also labeling vascular elements within the membrane. Notably, RPE cells within the lesion do not display any distinct polar labeling of the apical or basolateral sides with either lectin (Fig.1, arrows).

Both lectins showed strong labeling of choroidal blood vessels, especially the choriocapillaris. In order to more clearly determine the degree to which labeling was on the endothelial cell surface or on the closely apposed basal lamina, confocal microscopy was used to elucidate the structures that contain sialic acids that react with EBL and MAL-II. Co-labeling of each lectin with a marker of endothelial cell membranes (anti-CD34) and extracellular matrix (anti-Collagen IV) show that the labeling pattern of both EBL and MAL-II colocalizes most strongly with CD34 (Fig. 2). Some colocalization of both lectins can be seen with anti-Collagen IV, and in the healthy control this is stronger with EBL than MAL-II. No significant differences can be observed in the binding patterns between the control and early AMD maculae in the choriocapillaris. Of note, EBL and MAL-II labeling in the BLamD is consistent with the patchy and diffuse labeling observed at lower magnification.

**Figure 2.**
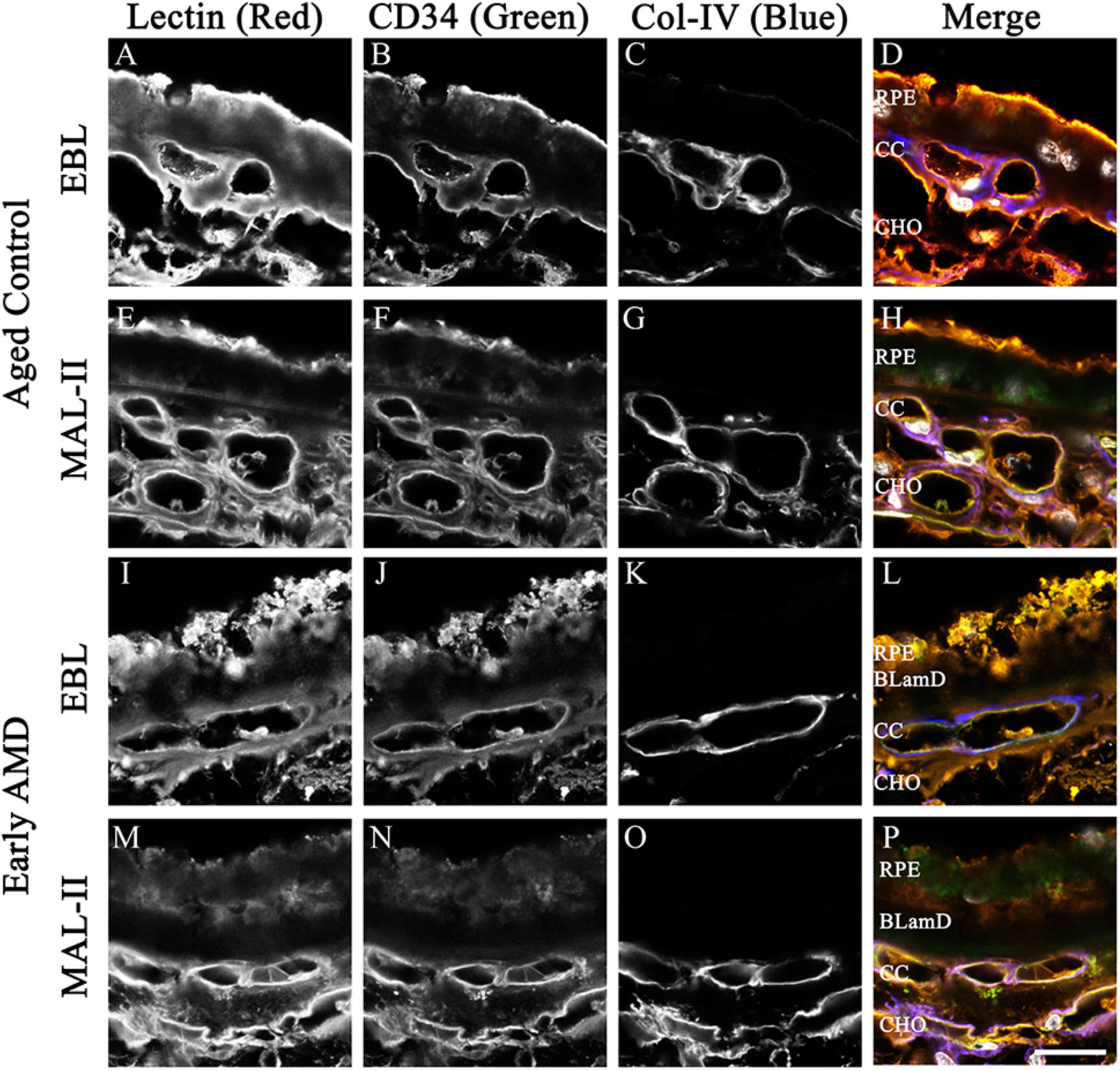
Colocalization of EBL or MAL-II with CD34 and Collagen-IV. Confocal micrographs of healthy (A-H) or early AMD (I-P) human choroids labeled with EBL (A,I) or MAL-II (E,M), anti-CD34 (B,F,J,N), and anti-Col-IV (C,G,K,O), each shown in grayscale. Merged images display EBL or MAL-II in red, anti-CD34 in green, anti-Col-IV in blue, and DAPI in grey. RPE, retinal pigment epithelium; CC, choriocapillaris; CHO, outer choroid. Scale bar is 20 µm.

As discussed above, FH and the C5b-9 MAC are of particular interest in the pathogenesis of AMD. Co-labeling of sections with anti-FH and anti-C5b9 was conducted to investigate the relationship between these two proteins/protein complexes and different sialic acid moieties (Fig. 3). FH and C5b-9 are observed in both the control and early AMD maculae. In the control, FH shows some overlap with both EBL and MAL-II labeling, though there is more overlap with EBL than MAL-II, consistent with EBL’s localization to the perivascular matrix and MAL-II’s localization to the endothelium. There is partial but incomplete overlap of FH and MAC in the intercapillary pillars (e.g., Fig. 3B, C). There are patches of FH positivity between the choriocapillaris and Bruch’s membrane that are not positive for either lectin (Fig. 3D, H). While lectin penetration was weaker in BlamD of the confocal sections, the overall pattern was similar to what was observed by epifluorescence, with stronger EBL labeling compared to MAL-II, especially on the apical surface of the deposit, which colocalized with FH (Fig. 3L).

**Figure 3.**
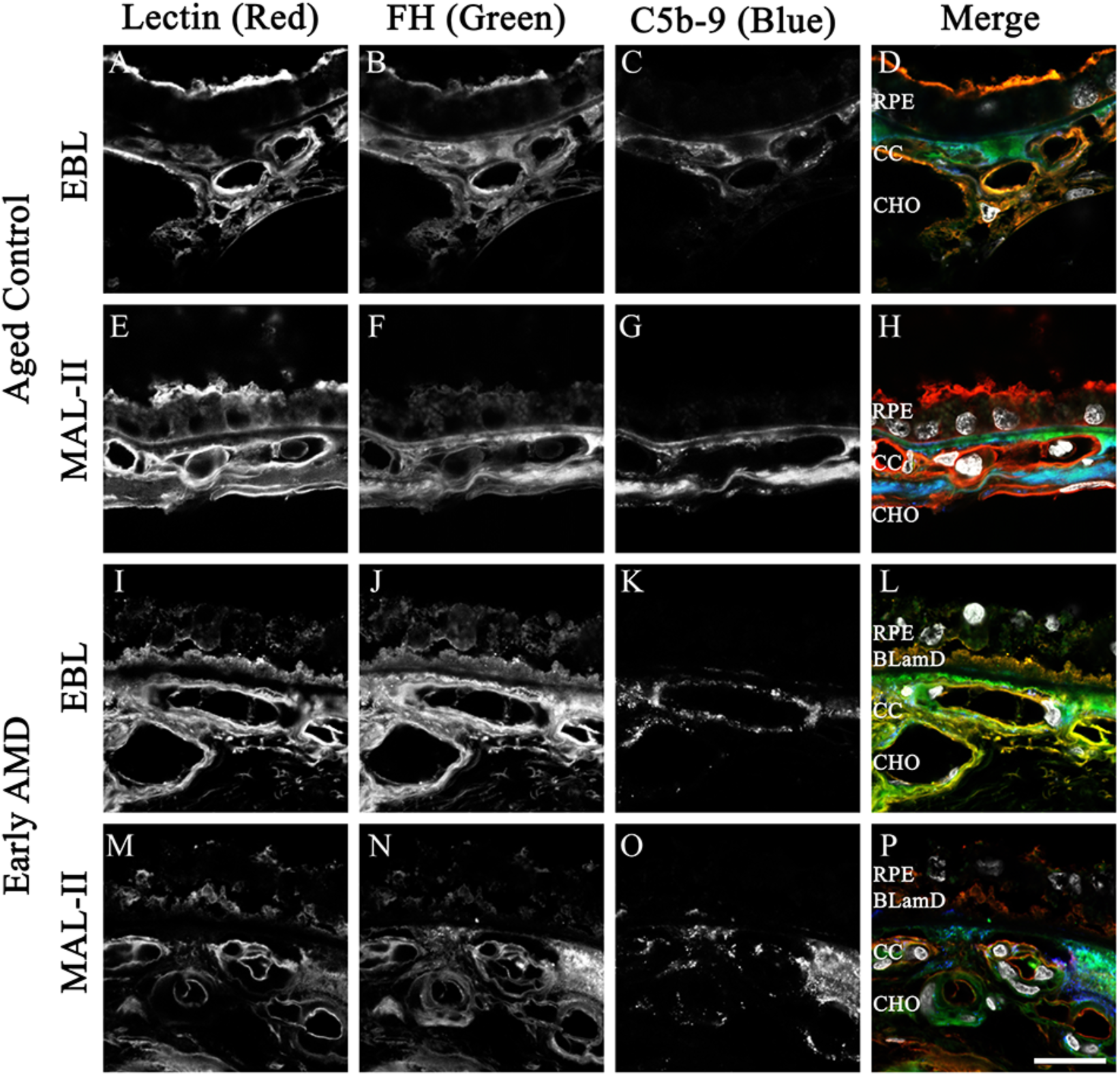
Colocalization of EBL or MAL-II with Factor H and C5b-9. Confocal micrographs of healthy (A-H) or early AMD (I-P) human choroids labeled with EBL (A,I) or MAL-II (E,M), and antibodies directed against Factor H (B,F,J,N), and C5b-9 (C,G,K,O), each shown in grayscale. Merged images display EBL or MAL-II in red, anti-Factor H in green, anti-C5b-9 in blue, and DAPI in grey. RPE, retinal pigment epithelium; CC, choriocapillaris; CHO, outer choroid. Scale bar is 20 µm.

To investigate the identity of molecules containing sialic acid moieties (glycoproteins or glycolipids), sections from two donors were treated with chloroform:methanol as described in the methods. Sudan Black B shows blue staining of Bruch’s membrane and BLamD in the untreated sections (Fig.4A, I), which is not present in the treated sections (Fig. 4E, M). In both donors, EBL and MAL-II labeling is decreased in the chloroform:methanol treated sections compared to untreated, especially in the choroidal stroma, capillaries, and larger vessels.

**Figure 4.**
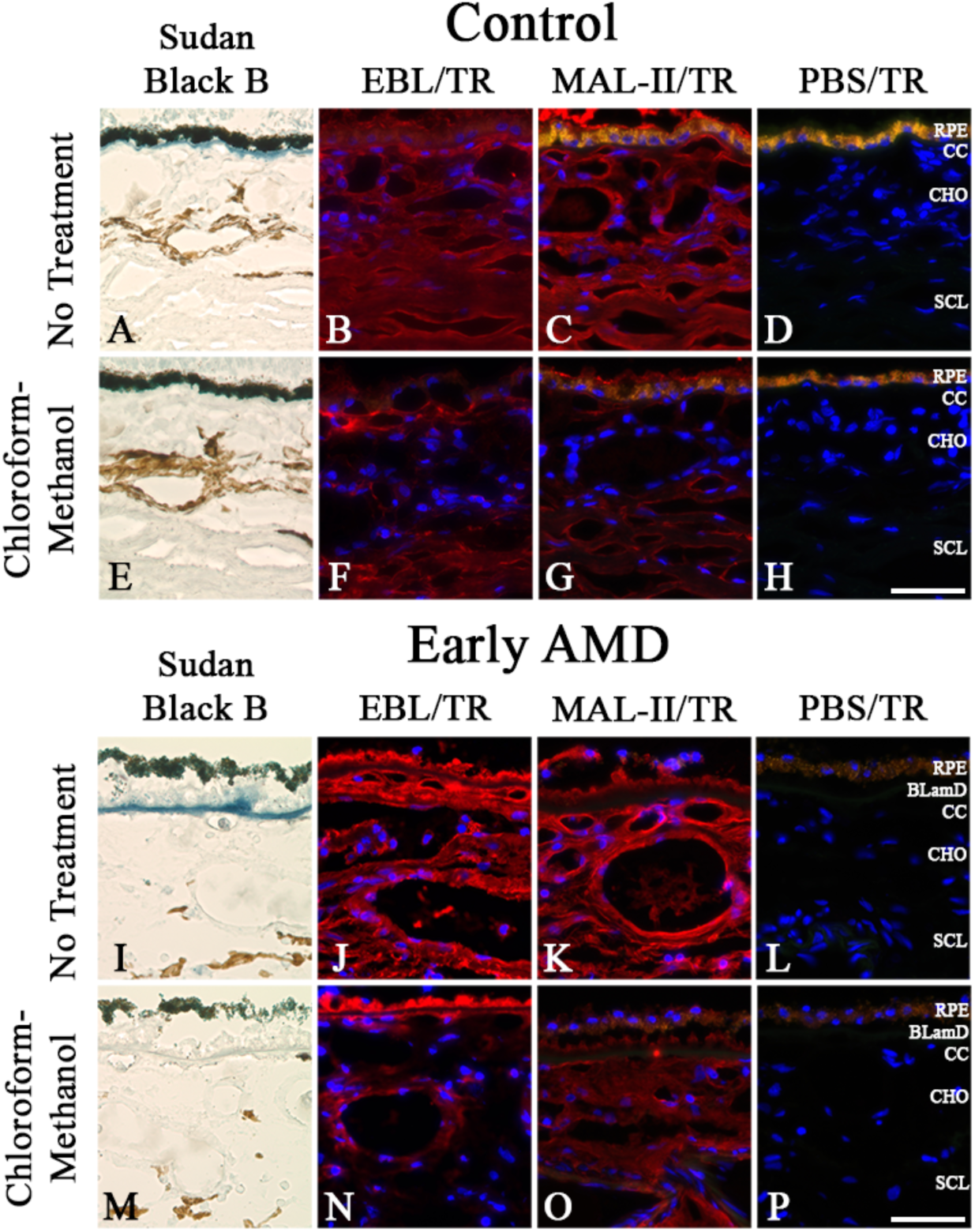
EBL and MAL-II labeling with and without chloroform-methanol treatment. Brightfield micrographs show Sudan Black B staining (A,E,I,M); fluorescence micrographs display EBL (B,F,J,N), MAL-II (C,G,K,O), or DAPI and Avidin Texas Red (TR) only (D,H,L,P). Panels A-H are from a healthy control, panels I-P are from a donor with early AMD and associated BLamD. RPE, retinal pigment epithelium; CC, choriocapillaris; CHO, outer choroid; SCL, sclera. Yellow-orange autofluorescence in control panels is due to RPE lipofuscin. Scale bar is 50 µm.

To further understand the composition of sialoglycoconjugates in the choroid, and to identify the penultimate glycans masked by sialic acid in different choroidal domains, human donor maculae were labeled with a battery of 37 lectins, with and without neuraminidase pre-treatment. The lectins employed react with mannose, fucose, galactose, GalNAc, GlcNAc, LacNAc, and other assorted complex glycan moieties. The change in labeling intensity between control and neuraminidase is displayed in Table 2. The lectins DBA, GSL-I, GSL-II displayed no labeling of the choroid or RPE, aside from very mild labeling of the choriocapillaris with GSL-I in two donors. Lectins that displayed decreased labeling with neuraminidase treatment included EBL and MAL-II as expected, as well as LEL, MAL-I, and WGA. Some lectins displayed variable changes with enzyme, both between donors and between compartments. An example of this is jacalin, which binds O-linked glycans. Jacalin showed increased labeling in Bruch’s membrane, stromal matrix, stromal cells, and BLamD, but decreased labeling in choriocapillaris and larger blood vessels. Other lectins showed strong increases in labeling following unmasking by neuraminidase treatment across multiple compartments, including BPL, ECL, PNA, SBA, SJA, and WFA.

Figure 5 shows examples of 4 lectins with and without neuraminidase treatment, including EBL, AAL, BPL, and ECL, in a donor with BLamD. EBL binds α-2,6 sialic acid and serves as a positive control. As expected, EBL signal is completely absent after neuraminidase treatment (Fig. 5A, B). AAL binds ɑ-Fucose and is an example of a lectin with mild increases in labeling intensity between control and enzyme treatment (Fig. 5C, D; note reactivity in BlamD). BPL and ECL bind β-galactose and LacNAc respectively, and labeling of both strongly increases following enzyme treatment (Fig. 5E-H; note labeling in choroidal stroma and BLamD). Minor labeling of choroidal vessels, stroma, and BLamD is present without the enzyme with both lectins. With neuraminidase treatment, there is intense labeling of these structures. This indicates that these anatomical features include glycoconjugates containing β-galactose and LacNAc, capped or obscured by terminal sialic acids.

**Figure 5.**
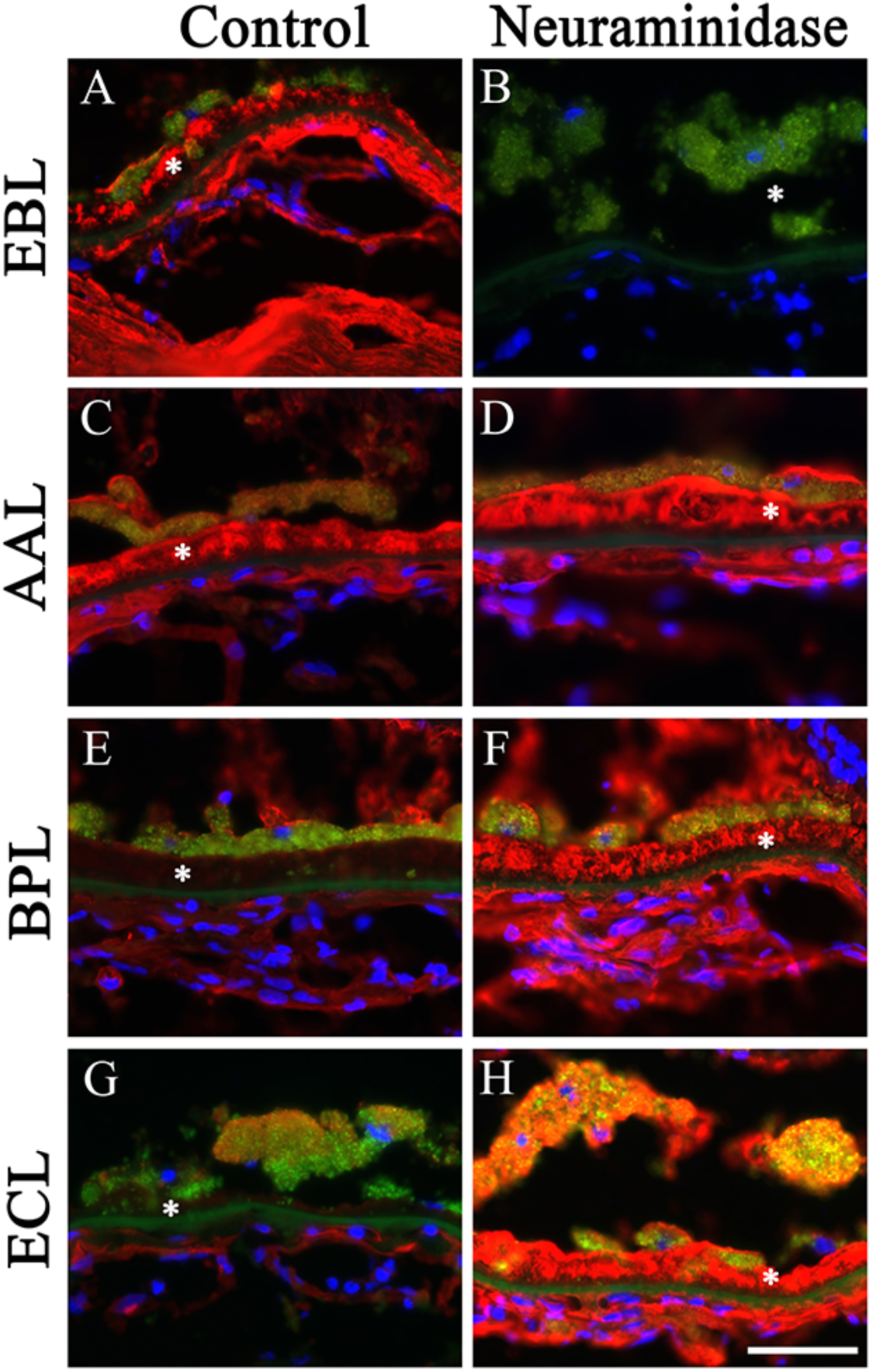
Labeling patterns of lectins with or without neuraminidase treatment. Fluorescence micrographs showing the labeling of EBL (A-B), AAL (C-D), BPL (E-F), and ECL (G-H). Asterisks (*) indicate BLamD. Scale bar is 50 µm.

## DISCUSSION

This study investigated the distribution and composition of sialoglycoconjugates in the choroid, in both healthy and disease states. Some research has been published regarding sialoglycoconjugates in retina (31) and glycoconjugates in drusen (25, 26), CNV lesions (27), and BLamD (29, 30), but little is known about sialoglycoconjugates in the healthy choroid and BLamD.

Our results show that both α-2,6 and α-2,3 sialic acids (shown by EBL and MAL-II respectively) are strongly associated with choroidal vasculature in both diseased and healthy states in different patterns. Both sialic acid forms are present in BLamD, whereas α-2,6 sialic acid is also prominent in the basal lamina and intercapillary pillars. It was noted that RPE cells within CNV lesions did not display typical polar labeling patterns with either EBL or MAL-II. This could be due to depolarization of the RPE within CNV lesions due to removal from their normal anatomical compartment and signaling partners.

Colocalization of EBL and MAL-II with markers for endothelial cell membranes (anti-CD34) and extracellular matrix (anti-Collagen IV) show that both are most strongly associated with the endothelial cell membranes, though EBL reactivity can also be observed in the extracellular matrix. Whereas previous studies have found that Factor H binds to α-2,3 sialic acid (34, 35), we found stronger colocalization with the α-2,6 sialic acid binding EBL. There are a few possible explanations for this finding. First, binding of endogenous FH to α-2,3 sialic acid could block MAL-II from binding those moieties, resulting in the observed lack of overlap between MAL-II and anti-FH. FH is itself a sialylated glycoprotein, with the majority of sialylation consisting of α-2,6 sialic acids (31); the colocalization of EBL and anti-FH in this study could be explained in part by EBL binding to α-2,6 sialic acid on the FH protein itself. Additionally, the colocalization for this study was conducted on two donors and without respect to *CFH* genotype; it is possible that different individuals display different patterns of binding and that FH binding in the choroid is complex. The C5b-9 MAC did not show clear association with either form of sialic acid. Decreases in EBL and MAL-II labeling after chloroform-methanol treatment suggest that a portion of the sialoglycoconjugates in the choroid are neutral glycolipids.

We further sought to identify penultimate carbohydrate moieties that may normally be masked by sialic acid. Lectin labeling after neuraminidase treatment showed a strong, consistent increase in labeling of β-galactose (BPL), LacNAc (ECL & SJA), Gal(β-1,3)GalNAc (PNA), and α- or β-GalNAc (SBA & WFA). Robust increases in labeling were noted in BlamD, pathologic deposits in macular degeneration that have a largely unknown composition.

This study provides insight into the sialoglycoconjugate profile of the choroid, including localization and partial molecular identities. This information increases our understanding of choroidal glycobiology, suggests avenues for future investigation into choroid and BlamD, and may provide a basis for targeting choroidal structures when designing treatments for AMD.

## ACKNOWLEDGEMENTS

The authors thank the eye donors and their families for their generous contributions to this research, as well as the Iowa Lions Eye Bank for their assistance in providing the tissue used for this study.

